# Contents of Consciousness Investigated as Integrated Information in Direct Human Brain Recordings

**DOI:** 10.1101/039032

**Authors:** Andrew M. Haun, Masafumi Oizumi, Christopher K. Kovach, Hiroto Kawasaki, Hiroyuki Oya, Matthew A. Howard, Ralph Adolphs, Naotsugu Tsuchiya

## Abstract

Integrated information theory postulates that the particular way stimuli appear when we consciously experience them arises from integrated information relationships across neural populations. We investigated if such equivalence holds by testing if similar/different percepts map onto similar/different information structures. We computed integrated information structure from intracranial EEGs recorded in 6 neurosurgical patients who had electrodes implanted over posterior cortices. During recording, we dissociated subjective percepts from physical inputs in three distinct stimulus paradigms (passive viewing, continuous flash suppression, and backward masking). Unsupervised classification showed that integrated information within stimulus-selective cortical regions classified visual experiences with significant accuracy (peaking on average around 64% classification accuracy). Classification by other relevant information theoretic measures such as mutual information and entropy was consistently poorer (56% and 54% accuracy). The findings argue that concepts from integrated information theory are empirically testable, promising a potential link between conscious experience and informational structures.

## Introduction

Conscious experience is intrinsic to the experiencing subject. This is most strikingly demonstrated during dreaming, hallucination, and imagination, where without any sensory inputs conscious experience emerge. For example, experiencing subjects usually cannot tell the difference between dreaming and normal experience (Nir & Tononi, 2010). Any moment of experience is also highly informative in the sense that it excludes a vast number of other possible experiences at that moment. Finally, experience is composed of many contents in a hierarchical structure that is irrevocably bound together into a holistic experience. These are the fundamental properties of consciousness: an intrinsic, integrated experience composed of various contents and that excludes other possible experiences. How can these fundamental properties of consciousness be supported by neuronal systems in the brain?

In the context of neuroscience, the concept of “information” is typically employed to characterize how much a neuron or neural population can tell an observer about presented stimuli or executed behaviors; receptive field mapping (Atick & Redlich, 1992; Seriès, Latham, & Pouget, 2004), object classification (Haxby et al., 2001), mind reading (Haynes & Rees, 2006; Kamitani & Tong, 2005), and response prediction can be all formalized in terms of mutual information *I*(*X;S*) or *I*(*X;B*), where *X, S* and *B* stand respectively for *states* of a neuronal system, *sensory* inputs, and *behavioral* outputs (Knill & Pouget, 2004; Quian Quiroga & Panzeri, 2009; Rieke, 1999). In other words, the mutual information quantifies how much uncertainty about stimuli or behaviors can be reduced if an *observer* knows the states of the neural system. Intrinsic information, a concept distinct from Shannon’s standard (extrinsic) “information”, is defined from an intrinsic perspective: how does a system’s state constrain its own future and past states (Oizumi, Amari, Yanagawa, Fujii, & Tsuchiya, 2016; Tononi, 2004)? It is the intrinsic information, better understood as *causal power* (Hoel, Albantakis, & Tononi, 2013; Oizumi, Albantakis, & Tononi, 2014), that is relevant for conscious experience, not the traditional notion of ‘extrinsic’ information. Consciousness exists even without stimulus, behavior, or an external observer: as a case in point, none of these are required for the rich experiences of dreams.

To ground the phenomenological notion of ‘integrated, intrinsic information’, the concept of *integrated information* has been rigorously developed within the framework of the *integrated information theory* (IIT) (Oizumi et al., 2014; Tononi, 2004) of conscious experience. However, the theoretical concepts of IIT are computationally difficult (or even intractable) for empirical observations and require approximated measures of integrated information. One such approximation can be derived via the mutual information between present and past states of a system (Barrett & Seth, 2011; Oizumi et al., 2016). The *entropy H(X(t))* of a system over an interval of time *t* quantifies the uncertainty of states *X* taken on by the system. The (auto-)mutual *information I*(*X*(*t*);*X*(*t* -*τ*)), can be understood as intrinsic information: how current states *X*(*t*) constrain possible past states *X*(*t-τ*), reducing the uncertainty of *X*. If we partition the system, estimate *I* only within the parts, and recombine these estimates, we discover whether or not some information is integrated across the whole system: the loss of information across the minimal cut of the system (information that cannot be confined to any partition) is the integrated information, Φ. Importantly, Φ is uniquely defined for all parts of a system. For example, the integrated information in a system of three elements, A, B, and C, is exhaustively characterized as the set {Φ(AB), Φ(AC), Φ(BC), Φ(ABC)}. We refer to this hierarchical pattern of integrated information as the *integrated information structure*.

The IIT proposes that consciousness (the intrinsic, integrated, and hierarchical phenomenon described in the first paragraph) is *identical* to the pattern of integrated information generated by the conscious system (i.e. the brain’s integrated information structure). Following from this, we hypothesized that the integrated information structure of a high-level visual area should closely reflect what subjects consciously perceive. We tested this hypothesis by measuring integrated information structures in human intracranial electrocorticography (ECoG) during various distinct perceptual contents. Perceptual contents were manipulated via stimulus paradigms including continuous flash suppression (CFS) (Tsuchiya & Koch, 2005) and backward masking (Breitmeyer & Ogmen, 2007), where physical stimulus and conscious perceptual contents are dissociable, and via unmasked stimulus paradigms. We extracted the intrinsic integrated information structure from small groups of ECoG channels, contingent on specific stimulus/percept conditions, and used these structures to classify perceptual contents. For comparison, we also extracted mutual information and entropy structures, which have related derivation but do not reflect information integration. We found that the integrated information structures mapped onto percepts better than related structures; in some subjects, the mapping was extremely precise and extended across multiple stimulus paradigms, consistent with the hypothesized equivalence between conscious contents and integrated information structure.

## Results

### Behavior

During ECoG recording, subjects participated in several visual tasks. We used both masked and unmasked stimulus paradigms (Supplemental Figure S1). In continuous flash suppression (CFS, Supplemental Figure S1A), a target face (neutral or fearful) and a flickering Mondrian mask were presented simultaneously into different eyes; the mask tended to suppress the target, especially when target contrast was low. In backward masking (BM, Supplemental Figure S1B), a face target (neutral, fearful or happy) was quickly followed by a noise mask, which tended to suppress visibility of the target when the mask-target delay was very brief. In the unmasked paradigms (UNM, one-back and fixation tasks, Supplemental Figure S1C), we showed upright/inverted neutral faces, Mondrians, tools, and houses; in different blocks, subjects were asked either to attend to stimulus category (one-back) or not to attend (fixation task). In CFS and BM, behavioral performance on a specific physical stimulus parameter allowed us to categorize individual experimental trials into those plausibly accompanied by specific conscious visual experiences of faces and other objects, while in UNM, we presumed that targets were visible so long as the subject continued to respond in the attention tasks (Supplemental Figure S1D). Behavioral performance is summarized in Supplemental Figure S2.

### Measuring Integrated Information

To estimate the integrated information structure in neural data, we used the time series derived from bi-polar re-referenced changes in field potential amplitudes recorded with subdural ECoG electrodes (Methods). We denote the (activity) state of a group of ECoG channels within a particular trial over time window *T* (from time t-T/2 to t+T/2) as the multivariate vector **X_t_**. We then estimate the covariance and cross-covariance matrices Σ(**X**_*t*_), Σ(**X_*t-τ*_**), and Σ(**X_*t*_**|**X_*t-τ*_**) with a lag parameter τ, for windows in individual trials (at this step, multiple trials’ worth of data can be combined by averaging over the covariance/cross-covariance matrices for each trial). Assuming Gaussian state distribution, we can then derive the entropy *H*(***X_t_***) and conditional entropy *H*(**X_t_**|**X_t-τ_**) (denoted below as *H_COND_*) over the time window.

The mutual information *I*(**X**_t_; **X_t-T_**) across the time lag τis the difference between the entropy and the conditional entropy, and can be understood as the information that the system possesses *about its own past*. It is important that for the mutual information, it does not matter whether parts of a system interact; even a collection of completely disconnected parts can have a large amount of information about its own past. Such a mere aggregate of independent parts should be quantified as zero with any measure of *integration*. A measure of integrated information should reflect the information generated by a *whole system*, above and beyond the sum of information generated by its parts (Tononi, 2004).

We estimate the integrated information, Φ^*^, as the difference between the mutual information *I*(**X**_t_; **X**_t-τ_) and the ‘mismatched decoding’ mutual information *I*^*^(**X**_t_; **X**_t-τ_) (Oizumi et al., 2016). *I*^*^ quantifies the mutual information generated by a model version of the analyzed system with *independent parts* (Latham & Nirenberg, 2005; Merhav, Kaplan, Lapidoth, & Shitz, 1994; Oizumi, Ishii, Ishibashi, Hosoya, & Okada, 2010) (See Methods as well). *I*^*^ is computed for the partition of the system that minimizes the difference between *I* and *I*^*^: the *minimum information partition* (MIP) (Balduzzi & Tononi, 2008). This minimal difference is information that cannot be isolated to parts of the system: the integrated information, Φ^*^ = *I* -*I*^*^ (for detailed derivation, see Methods and (Oizumi et al., 2016)).

### The Structure of Integrated Information

To mirror the nested, multi-order structure of perceptual experience, we measured Φ^*^ for every subsystem within a selected system of ECoG channels. The resulting ‘power set’ of Φ^*^ values describes a hierarchy of overlapping subsystems that may or may not integrate information within the specified system - we refer to this hierarchical set of integrated information values as the *<P* structure*. We measured these Φ^*^ structures at every location of a 4-channel searchlight, in each of six subjects, for a range of τ (time lag) values from 1.5 to 12 msec. All computations were performed over 200ms time windows (longer windows make for better covariance matrix estimates, while sacrificing temporal specificity) spaced 100ms apart over a period from 400ms before stimulus onset to 1000ms after onset. To select regions of interest (presented in Figures 1 through 4), we focused on maximae in evoked Φ^*^, i.e. the searchlight locations with maximal increase in Φ^*^ after stimulus onset, regardless of the stimulus category. Our rationale for this selection was theory-driven: we reasoned that where information integration is evoked by a stimulus, the *structure* of the information should naturally identify the percept. The procedure is described in detail in the Methods. Similar criteria for selecting a region of interest are discussed in the Methods and illustrated in Supplemental Figure S4.

**Figure 1.**
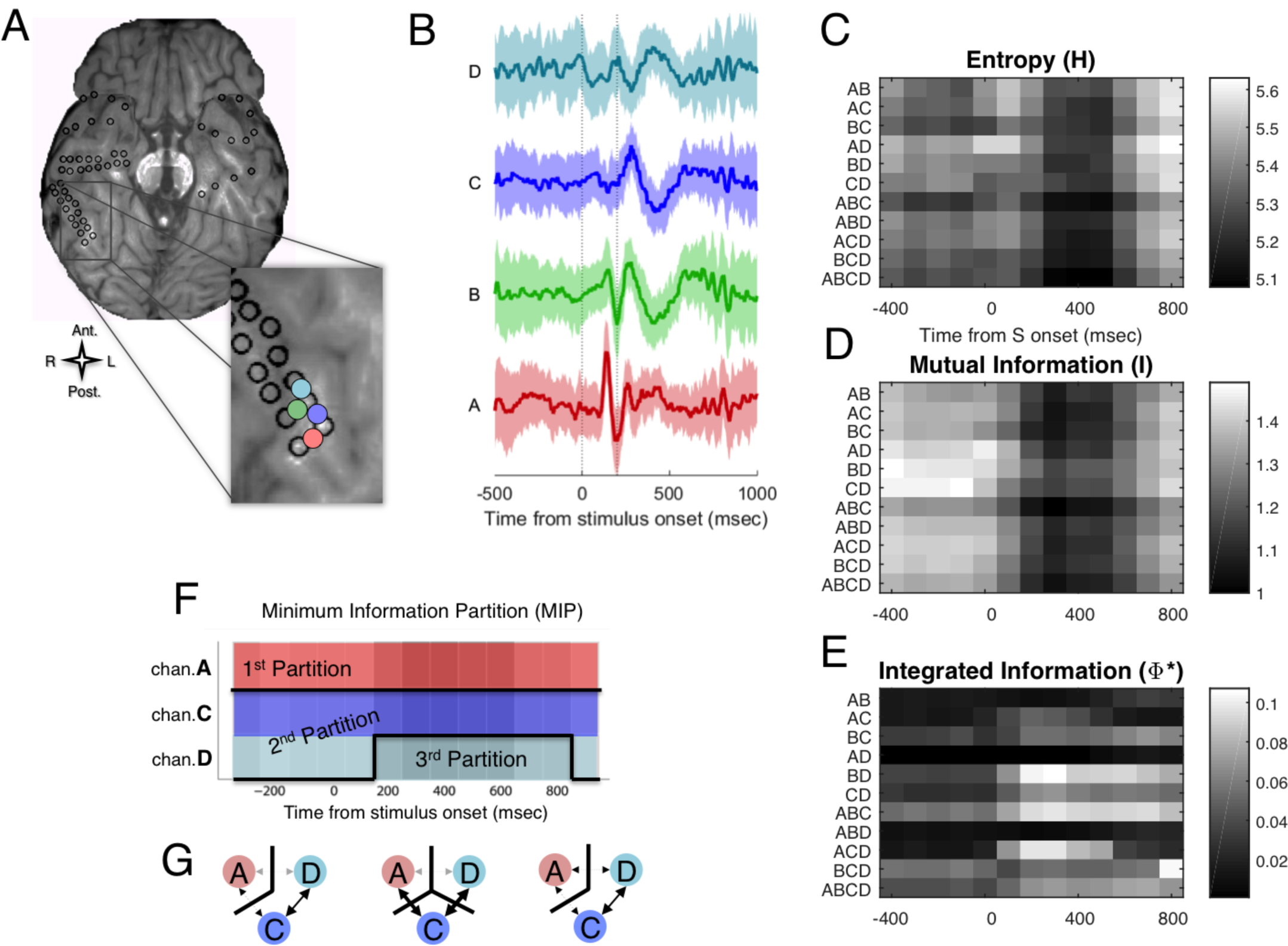
The procedure for estimation of integrated information structure in ECoG data. A) ECoG recording. Black rings mark the location of electrodes; we used bipolar re-referenced channels between each local pair of electrodes. Four of these are marked in color. B) Means (thick lines) and standard deviations (shades) of the field potentials for the four channels marked in A, from 500ms before to 1000ms after a face stimulus onset. Here we averaged intervals over 45 trials in the CFS experiment where the high-contrast face target was reported as highly visible by Subject 153. C-E) Time courses of the entropy *H*(*C*), mutual information *I*(*D*) and integrated information Φ^*^(E) for each of the 11 subsystems. Each subsystem is a subset of the channels in the system ABCD. Values were estimated over a time window T=200ms and τ = 3ms, and plotted by aligning its center with the time point on the x-axis. Entropy and the mutual information are proportional to number of channels, so we have plotted H and I divided by the number of channels for each subsystem, to emphasize the dynamics over all subsystems. The dynamics of Φ^*^ are highly idiosyncratic. Note the increase in Φ^*^ for subsystems BD and ACD, after the stimulus onset. The increase in ACD’s Φ^*^ is accompanied by change in its minimum information partition (MIP) (F and G): subsystem ACD switches from a bipartite to a tripartite MIP when a face is seen, accompanied by the increase in Φ^*^ magnitude (indicated by the darker colors).

We first present a detailed analysis of integrated information structure in a single subject (S153), in a group of channels located over the right fusiform gyrus (FG), a region that has a known association with conscious perception of faces (Baroni et al., 2016; Parvizi et al., 2012; Puce, Allison, & McCarthy, 1999; Rangarajan et al., 2014; Tong, Nakayama, Vaughan, & Kanwisher, 1998). The process is illustrated in Figure 1 for a system of four channels (Figure 1A) - these four channels (with τ=3msec) contained the highest evoked Φ^*^ of any searchlight location in this subject. Figure 1B plots the average time course of raw bipolar re-referenced voltages (no baseline correction) during trials where subjects consciously saw (visibility rating of 4) high-contrast faces in the CFS task. We computed Φ^*^ for *every combination of channels* within this system: for a system of 4 channels this yields six 2^nd^-order subsystems, four 3^rd^-order subsystems, and a single 4^th^-order subsystem. Panels **1C-E** show the time courses of the quantities that underlie Φ^*^, computed for each of 11 subsystems. The most general quantity is the entropy *H* (Figure 1C). From the entropy *H* we subtract the conditional entropy *H_COND_*, yielding the mutual information *I* (Figure 1D). Next we identify the MIP for each subsystem, which identifies the weakest link that minimizes the difference between *I* and *I\** (the difference is subject to a normalization term; see *Methods;* also note that for a 2^nd^-order subsystem, the only possible partition is the MIP). The integrated information is evaluated at the MIP: Φ^*^ = *I* - *I\** (Figure 1E).

Most subsystems show a similar time course for *H* and *I* (in the example system of Figure 1 as well as most other observed systems), with entropy and mutual information decreasing a few hundred milliseconds after stimulus onset. In contrast, Φ^*^ has an idiosyncratic time course, strongly dependent on the specific subsystem. For most subsystems, Φ^*^ remains near zero regardless of the visual stimulus; for others, it may start high and drop after a particular stimulus is presented (e.g. subsystem BCD in Figure 1E); and for other subsystems Φ^*^ starts low and increases after stimulus presentation (e.g., subsystems BD and ACD in Figure 1E). A further point of interest around the Φ^*^ dynamics is the internal structure of each computation: the *MIP*. Figure 1F shows how the MIP changes over time for a particular subsystem ACD, resting in a bipartite state (A vs CD) before stimulus onset, and switching to a tripartite state (A vs C vs D) after stimulus onset. This change in partition structure accompanies/underlies ACD’s increased integrated information upon the onset of stimulus onset.

### Graphing *Φ^*^* Structure

The complete set of subsystems within a specified system constitutes a partially-ordered set that can be visualized with a Hasse graph (Figure 2A). In the graph (whose x- and y-coordinates are arbitrary), we place the highest 4^th^-order subsystem ABCD at the origin (the red colored node), surrounded by the 3^rd^-order subsystems in black, which are further surrounded by the 2^nd^-order subsystems in blue. The edges represent the simplest possible transition through a ‘subsystem space’: one subsystem transitions to another by adding or subtracting one channel. For example, subsystem ACD, the 3-order node featured in Figure 2F, is connected to subsystems AC, CD, AD via edges as well as ABCD. The 3D rendering of the structure is easily appreciated in Supplementary Movie 1.

**Figure 2.**
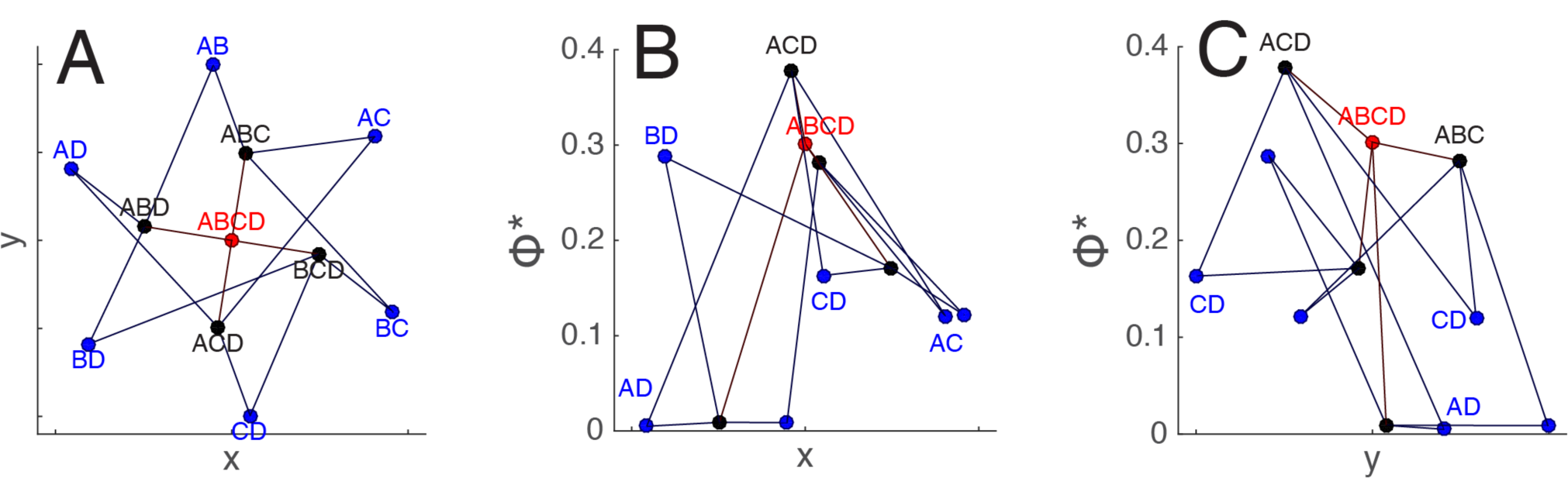
Integrated information structure presented on a Hasse graph. See also Supplementary Movie 1. A) The graph for all subsystems in the system ABCD evaluated in Figure 1 in the interval 200-400 ms. The *x* and *y* coordinates here are assigned for visualization purposes. Each node in the graph is one of the 11 subsystems in the system ABCD. The edges of the graph represent addition or subtraction of a single channel from a subsystem. The color of each node (when visible) represents the order of the subsystem: blue nodes are 2^nd^-order subsystems, black nodes are 3^rd^-order subsystems, and the red node is the ‘top’ 4^th^-order subsystem. **B**) The same graph viewed against the y-axis, with the magnitude of Φ^*^ represented on the vertical (Z) axis. Subsystem ACD is labeled in black – adding or subtracting channels to this subsystem only reduces Φ^*^. All the other subsystems in this system integrate less information than subsystem ACD, including the ‘enveloping’ higher-order system ABCD. **C**) The same graph viewed against the x-axis.

The 4-channel Φ^*^ structure in Figure 2 is suggestive of a highly nonlinear and nonmonotonic changes in the shape of the information integration structure. Here the 3rd-order subsystem ACD attains the highest Φ^*^ of all 11 possible subsystems. The graph illustrates how subtracting from *or* adding to a subsystem can reduce the integrated information. The Φ^*^ of subsystem ABCD is *less* than that of ACD because there is a relatively weak interaction between B and ACD. In this case, the *minimum information partition* (MIP) is between B and ACD.

Figure 3 shows Φ^*^ structures computed over the same time period (200 to 400msec after the stimulus onset) in the same group of channels as in Figures 1 and 2. Figures 3A-C present Φ^*^ structures obtained in three separate experiments: the unmasked one-back task, backward masking (BM) and continuous flash suppression (CFS) (For comparisons of these structures, see Supplementary Movie 2 to view Φ^*^ structures in a 3 dimensional perspective). Φ^*^ structures in the left two columns are constructed from trials where the subject clearly perceived a face (81 trials of unmasked upright or inverted faces in the one-back task, 68 trials with correct in both location and emotion discriminations in BM, and 94 trials with high visibility ratings in CFS, with hit/miss counts identified in the inset text), whereas the right two columns are not (38 Mondrian trials in the one-back task, 28 trials with incorrect in location discrimination in the BM, and 95 trials with low visibility ratings in the CFS). In this example, Φ^*^ structures in trials with clearer face percepts (the left columns) have higher, tilted shapes, while non-face percepts correspond to flatter shapes. The trends are carried by differential behaviors in some subsystems; for example, subsystem ACD (marked with the orange circles) attained high values of Φ^*^ only when a face was visible (left columns), while subsystem ABC (marked with blue circles), was integrated regardless of perceptual state.

**Figure 3.**
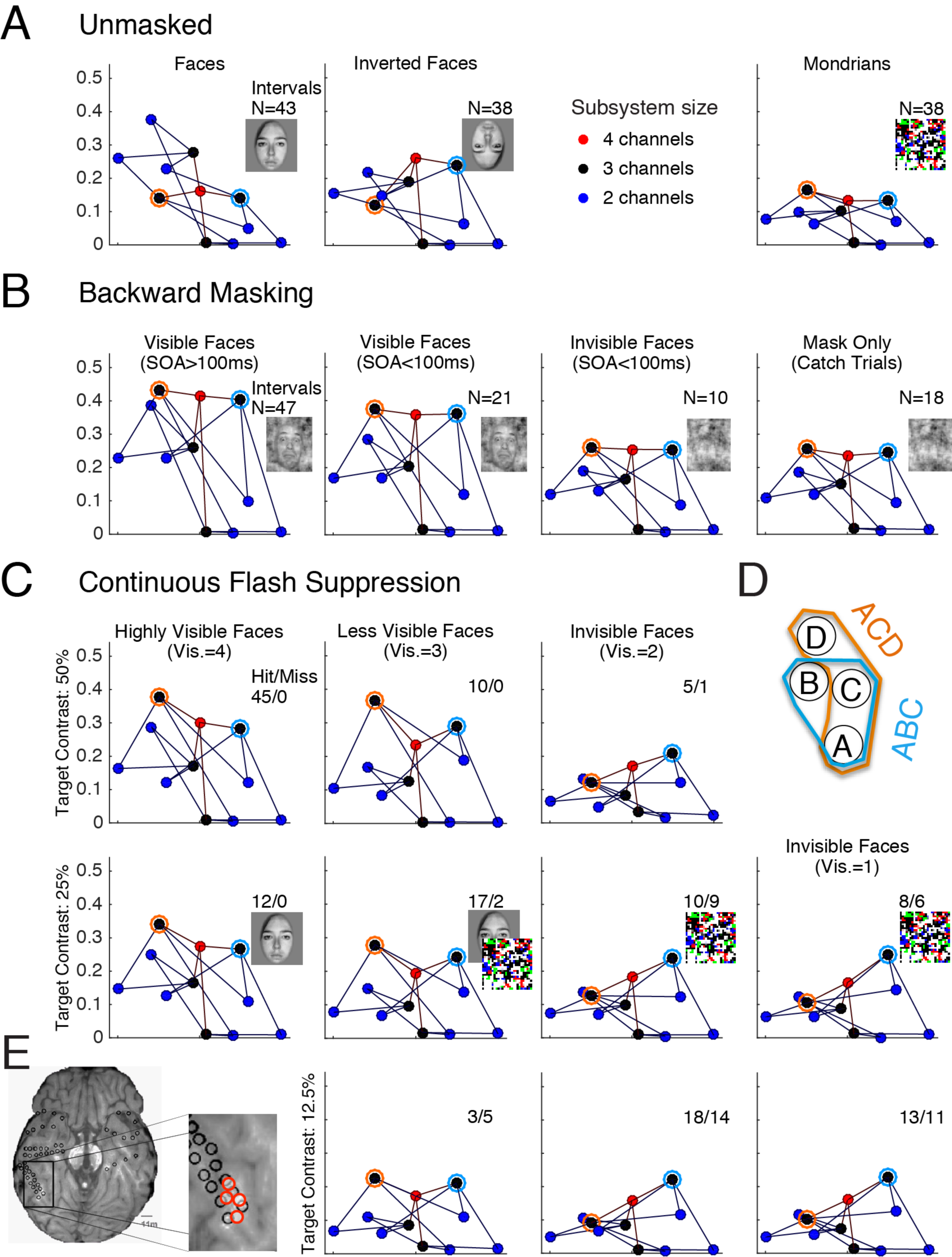
A-C) Graphs of Φ^*^ structure in the interval 200-400msec after stimulus onset, in S153’s right fusiform gyrus (location specified in panel **E**), in multiple percept/stimulus conditions in three stimulus paradigms. Marker colors mean the same thing as in Figure 2. The same channel system in Subject 153 is analyzed here as in Figures 1 and 2. Two prominent subsystems, ABC and ACD, are marked by the blue and orange circles in each graph – these are indicated in D. A) Φ^*^ structures for trials with the three unmasked stimuli in the one-back experiment: Faces, Inverted Faces, and Mondrians. **B**) Φ^*^ structures generated in the four trial types in the backward masking experiment: Visible Face trials here are those where the subject correctly localized (4AFC) and identified emotion (3AFC) of the masked face, shown separately for long and short SOA trials; Invisible Face trials are those short SOA trials where the subject was incorrect for localization; and ‘Mask Only’ trials were catch trials where no faces were presented. **C**) Φ^*^ structures from the CFS experiment. Within each row, physical structure of the stimuli, including contrast of the target face, was identical. Columns indicate the reported visibility of the face target. Hit/miss ratios are shown in each panel. The inset images roughly depict *what was perceived* in the corresponding intervals. To compute Φ^*^ structure in each panel, we used the number of trials available in the condition (N or Hit+Miss), as indicated by the inset numbers. See Supplementary Movie 2 for comparison of visible face trials in CFS and BM with invisible trials in CFS.

The overall shapes of the structures are similar within the columns, implying that these Φ^*^ structures reflect conscious perception of faces, regardless of the task instructions and low-level visual stimulus properties. This is consistent with a proposed equivalence between conscious perception of faces and the Φ^*^ structures, which are invariant to task or stimulus details (Aru, Bachmann, Singer, & Melloni, 2012; de Graaf, Hsieh, & Sack, 2012).

### Natural classification of conscious contents

To objectively assess the link between Φ^*^ structure and perceptual contents, we employed classification analyses based in the notion of representational similarity (Kriegeskorte, Mur, & Bandettini, 2008). We computed <D* structure over bins of 3 trials each matched for percept and stimulus categories (e.g., high-contrast CFS faces reported as visible). <D* structure can be characterized in at least two ways. First, it can be represented as a vector of Φ^*^ magnitudes (|Φ^*^|) for each of N subsystems (with 4 channels, N=11); we refer to this as the |Φ^*^| representation. Second, <D* structure can be represented as the pattern of MIPs over N subsystems (e.g., tri-partition of subsystem ABC (A vs B vs C), or bipartition (AB vs C)); we refer to this as the MIP representation. Here, we measured similarity of the <D* structure in these two xsdomains separately (Methods).

To evaluate the mapping between percept and structure, we constructed matrices representing dissimilarity of Φ^*^ structure between every pairing of trial bins (1 minus the Pearson correlation coefficient (Kriegeskorte et al., 2008) for |Φ^*^| dissimilarity, and a simple matching measure for MIP dissimilarity as described in *Methods)*. For comparison, we measured dissimilarity for entropy (*H*) and mutual information (*I*) structures, which are obtained in the course of the Φ^*^ computation. We then projected the dissimilarity matrices into multi-dimensional scaling (MDS) coordinates, and used these coordinates to measure percept classification accuracy as the area under the receiver operator characteristic curve (AUC) for each structure type, at each searchlight location (see Methods for details). Importantly, there was no training step to optimize weights for classification: the raw values extracted from the structures served *directly* as unweighted feature vectors. The searchlight AUC results are summarized in Figure 4a, showing the likelihood of each structure type given a specified AUC, for each of six subjects; while very high AUC values are rare (only a few searchlight locations in each subject), when AUC *is* high it is with |Φ^*^| or MIP structures, and never with *I* or *H* structures. Φ^*^ structures naturally classify the contents of consciousness. Thus our main results that |Φ^*^| or MIP classifies better than H or I do not crucially depend on the exact channel selections, at least when we analyze the high-level visual areas in this study.

**Figure 4.**
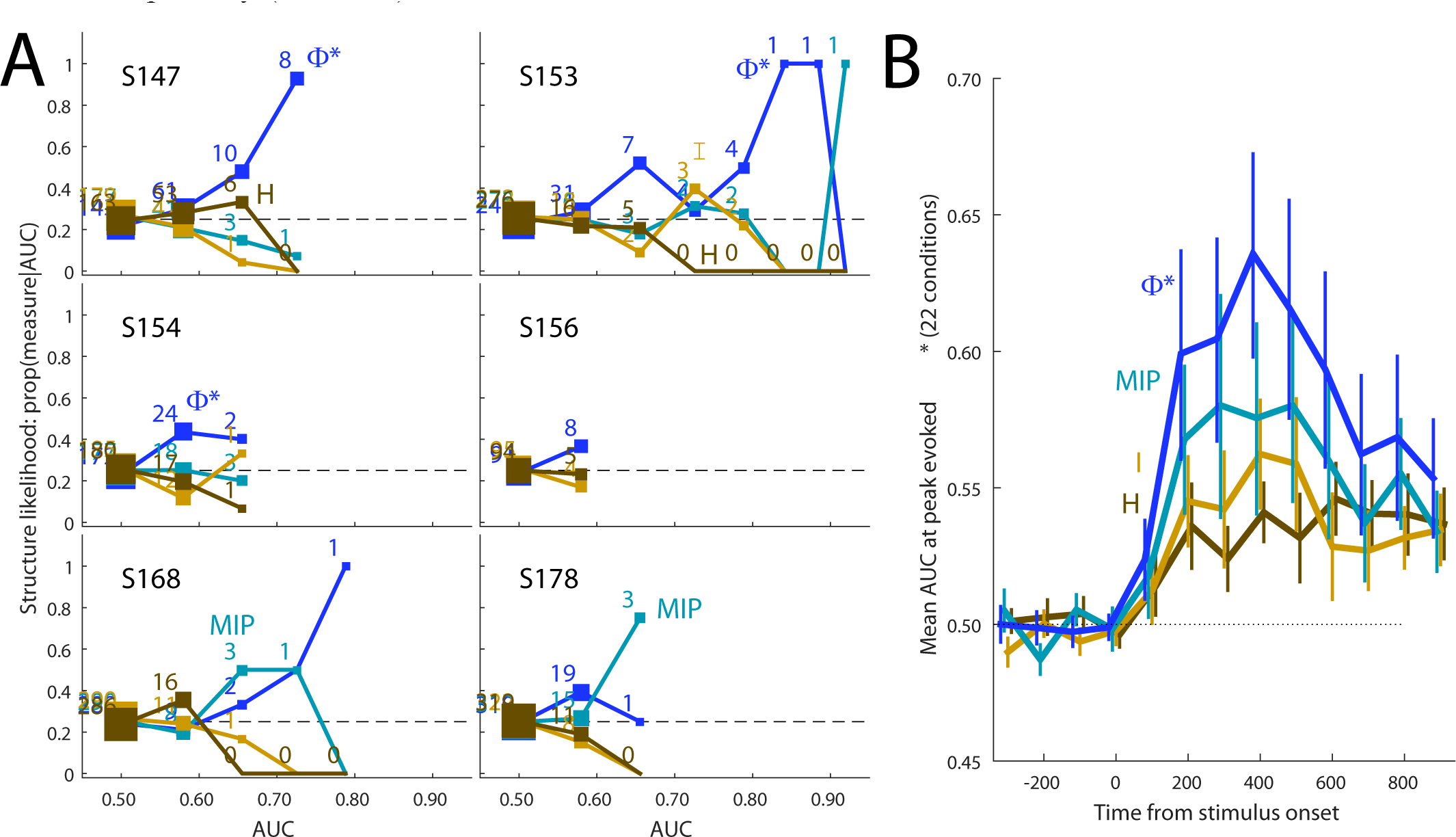
Percept classification accuracy for different information structure types. **A**) The likelihood of different structure types given specific classification accuracies (AUC), for each subject, combined over all searchlight locations in all stimulus paradigms. Likelihood for each structure type is the incidence of that type at the given AUC, divided by the total number of structures observed at the given AUC. When classification accuracy is high (value on the x-axis), it is usually obtained (with high likelihood) with Φ^*^ or MIP structures. The numbers by each marker indicate the number of observations of each AUC value, for each structure type. Chance AUC is 0.5 (see Methods); chance likelihood, since there are four structure types (O, MIP, I, H) is 0.25. **B**) Average timecourse of discriminability over 22 conditions (combinations of six subjects, three possible tasks, and either lateral or ventral electrode implantations). |Φ^*^| and MIP structures classify percept category better than mutual information and entropy (I and H). Error bars are the standard error of the mean of the z-transformed AUC scores.

What might mark these regions of strong Φ^*^-percept mapping? We reasoned that where information is integrated in response to a stimulus, its structure should map closely to the reported experience. In each of 22 distinct conditions (six subjects, most with lateral and ventral ECoG installations, each of whom performed at least one of the visual tasks: not every subject completed every task, and not all had ventral electrodes; see Table 1 in Methods) we selected the searchlight location that generated the largest evoked Φ^*^ regardless of stimulus/percept condition (the same criterion by which the location presented in Figures 1-3 was selected). Figure 4b shows the AUC timecourse for each measure, averaged over the 22 conditions; average classification is best with |Φ^*^| and MIP structures. We used the post-stimulus onset AUC scores (z-transformed) over all conditions (combinations of 6 subjects, 3 tasks, and two types of electrode installation; see methods) *and* using all structure types (Φ^*^, MIP, I, and H) as dependent measures in an ANOVA, with structure type, subject ID, task, and installation type as fixed factors and post-stimulus time point as a covariate. There was a main effect of structure type (F(3,703)=13.290, p<0.001), with significant (Bonferroni-corrected) post-hoc pairwise differences between AUC derived from Φ^*^ versus *I* structures (p<.001), Φ^*^ versus *H* structures (p<.001), and Φ^*^ versus MIP structures (p=.006) (these comparisons reflecting differences between the post-stimulus time courses in Figure 4b). MIP-derived AUC was also significantly different from H-based AUC (p=.006). There were significant main effects of subject ID (F(5,703)=89.214, p<.001), task (F(2,703)=52.281, p<.001), and installation type (lateral/ventral; F(1,703)=68.851, p<.001). All factorial interactions were significant. Other (less theoretically-motivated) selection criteria that are aimed simply at identifying percept-informative cortical regions yield qualitatively similar results (Supplemental Figure S4).

## Discussion

According to IIT, conscious experience is identical to a structure of intrinsic integrated information (Tononi, 2004); so, features of integrated information structure observed in a brain *should* correspond to features of the subjective experience of that brain. In this paper, we presented a tool - the integrated information measure Φ^*^ - by which such equivalence can be investigated. We found that integrated information structure in human visual cortex naturally classified psychophysically determined perceptual contents, across task instructions and in conditions where physical stimuli are dissociated from perceptual experience. Importantly, Φ^*^ structure maps more closely to perceptual states than do entropy and mutual information structures, despite their overlapping derivation. The advantage of Φ^*^ over entropy and mutual information is likely due to Φ^*^’s isolation of *integration*. Entropy quantifies only the instantaneous uncertainty of the neural states, understandable as equal-time interactions (Oizumi, Tsuchiya, & Amari, 2015). Mutual information quantifies time-lagged interactions in these distributions, i.e., how past states affect future states, including both within and between channels (Figure 1B). Φ^*^ also quantifies time-lagged interactions but only those that tie groups of channels together. Thus, our results can be summarized as showing that neither the structure of equal-time interactions (H) or of time-lagged self-interactions (I) contain as much information about conscious percepts as do time-lagged cross-interactions (Φ^*^). Thus, our results suggests a critical role of time-lagged interactions across the system, or integration, of neuronal activity for understanding how conscious phenomenology corresponds to neural systems.

We observed integrated information structure whose emergence tracked closely with the experimentally controlled contents of consciousness. In the notable case of Subject 153, we observed a similar structure generated in different stimulus paradigms when the subject experienced a ‘face’ percept. The elements generating the structure - channels of neural activity in the right fusiform gyrus - have long been associated with face percepts on the basis of patterns of responsivity and perceptual consequence of electrical stimulation of the area (Grill-Spector, Knouf, & Kanwisher, 2004; Haxby, Hoffman, & Gobbini, 2000; Parvizi et al., 2012; Tong et al., 1998). Here we have demonstrated that the neural activity in this region that reflects face *awareness* is irreducibly integrated across multiple hierarchical levels. Generalizing this finding, we have demonstrated in a number of subjects, stimulus paradigms, and cortical regions, that *how the information is integrated* (the Φ^*^ structure) indeed reflects the nature of the percept evoked by a stimulus. The caveat, typical of ECoG recording in human patients, is that our data are influenced by large variability in electrode implantation locations and in each subject’s behavioral performance, which likely contributed to the variance in the quality of classification across condition (e.g., the outstanding result from patient 153, who was cognitively alert and whose electrodes seem to have been in exactly the right place). Given the difficulty in conducting this type of experiment in epilepsy patients, it will be important to continue accumulating data to determine how the conclusions from this study extend to other patient samples.

By using a measure of integrated information, we have applied several new concepts to the analysis of neural data. However, for our purposes it is the concept of integrated information *structure* that is most crucial, as this concept ties into the hierarchical, compositional nature of the contents of consciousness. The integrated information theory provides us with a detailed prediction that <D* structure should have a close connection to specific conscious contents (Balduzzi & Tononi, 2009; Oizumi et al., 2014). A conscious experience is *intrinsic,* existing for itself, not some outside observer - Φ^*^ measures how a system specifies its *own* state, not an external state; a conscious experience is *integrated,* irreducible, more than the sum of its parts - Φ^*^ measures how the *system as a whole* specifies its own state, above and beyond its elements; and a conscious experience is *compositional,* just like the hierarchical Φ^*^ structure. Based on these theoretical parallels, there is strong a priori reason to expect Φ^*^ structures should closely correlate with perceptual experience. This type of theoretical background is lacking in many investigations of the neural correlates of consciousness; rather than the traditional search for correlates, our study can be seen as a test of a theory.

Our results should be seen as a step in the direction of a new empirical research framework in consciousness science, where the degree of equivalence is assessed between the domain of conscious experience and the domain of a theoretical construct, with empirical data as a bridge between domains (Tsuchiya, Taguchi, & Saigo, 2016) If the equivalence holds, similar experiences should map onto similar Φ^*^ structures (or other, further developed constructs), and different experiences should translate to different structures, as we saw in this paper. Looking forward, the IIT would say that for local O structure to be part of conscious experience, it must be a substructure of a vastly larger ‘main complex’ extending across multiple cortical regions. Measuring this superstructure in a paradigm where local perceptual structure can be simultaneously resolved seems a formidable task, but the difficulties may lie more in data type and availability than in computational complexity (as our results suggest). These difficulties will be addressed by future experiments and by further theoretical developments. With the methodological framework we provided here, it will be possible to test the theory with other datasets in other experimental manipulations combined with neural recording and stimulation, approaching ever closer to establishing the link between the mental and the physical worlds.

## Methods

### Subjects

We analyzed ECoG recordings obtained in six patients undergoing video EEG monitoring with intracranial electrodes. All patients had ‘grid’ arrays installed over the left (N=4) or right (N=2) lateral temporal lobes, and five also had two or more ‘strip’ arrays installed ventrally on the same side. Patients also had frontal and deep electrodes, which we did not include in our analyses. Recording was not performed within 48 hours of major seizures (Lachaux, Axmacher, Mormann, Halgren, & Crone, 2012). Subjects, experimental paradigms, and the cortical area that received electrode implantation for all 22 conditions considered. For the analysis reported in Figure 4, we regarded each row in Table 1 as an independent condition.

**Table 1.**
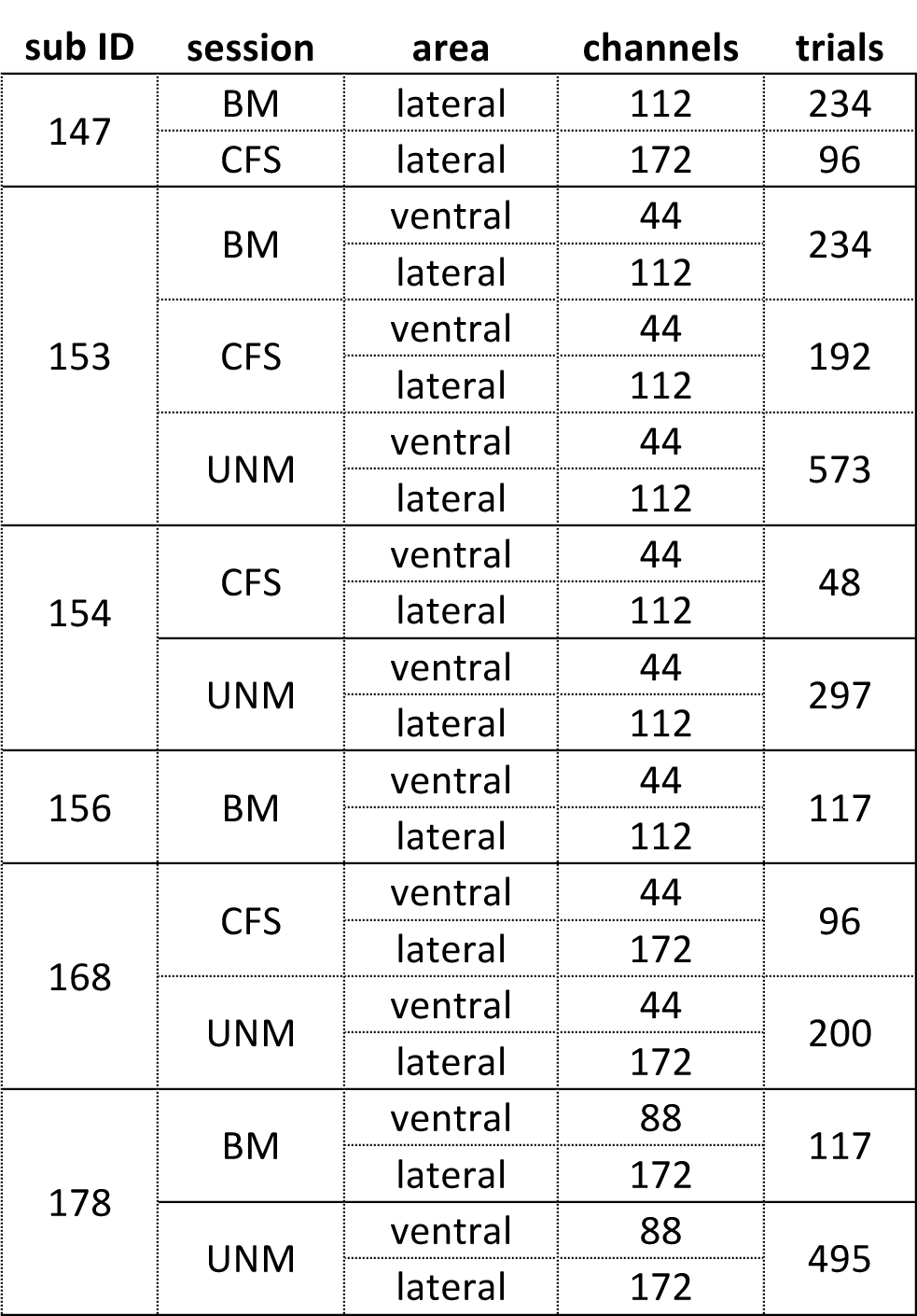

## Psychophysics

Subjects performed psychophysics experiments while outfitted with the ECoG arrays. All subjects were naïve to the tasks, and generally received a block or two of practice trials before data collection began. Completed trial counts for all included subjects are listed in Table 1.

Continuous flash suppression (CFS) is a dichoptic masking paradigm. In our experiment, a target face stimulus is presented to one eye, while distinct patterns of colorful Mondrian (shape noise) images are continuously flashed to the corresponding position in the other eye (Tsuchiya & Koch, 2005). In each trial, two temporal intervals were presented (Figure S1a). Each interval lasted 200msec, and the two intervals were separated by a random duration between 900 and 1100 msec. In one interval, the target face was presented to one eye; in the other interval, the blank gray field was presented. In both intervals, three distinct patterns of Mondrians were presented (each 66 ms, at 15Hz) to the other eye. Dichoptic view was achieved through a custom-made mirror scope consisting of 4 mirrors.

After the two intervals, the subject was asked to report which interval contained the target face, and then to report the subjective visibility of the target (a 4-point scale ranging from ‘clearly visible’ to ‘invisible’) (Rams0y & Overgaard, 2004). Three target contrasts (either 50%, 25%, and 12.5% or 100%, 50%, and 25%, depending on the subject) were interleaved from trial to trial. As a screening step (to ensure that included subjects understand the task and responded properly), we included only data from experimental sessions whose objective 2AFC hit rate increased with target contrast, *and* increased with reported visibility. If these criteria were met, we treated visibility levels where hit rate was near chance (50%) as “invisible targets”, and higher visibility levels as “visible targets”. For all included CFS subjects, a division of trials that separate trials with visibility ratings of 3 (mostly visible) or 4 (clearly visible) and ratings of 1 (complete guess) and 2 (nearly invisible) satisfied the above criteria. Summary measures of CFS performance for the included subjects are shown in Figure S2a.

In the backward masking (BM) paradigm, a target face stimulus, whose emotional expression was either happy, angry, or neutral, was briefly flashed (17 msec) at one of four visual field locations (upper-left, upper-right, lower-left, lower-right), which was immediately replaced by a gray blank screen. The face was placed within an oval shaped mask within the 1/f noise. The other three quadrants contained 1/f noise only. After a variable stimulus onset asynchrony (SOA), the stimulus array that included the target face was replaced with 1/f noise in all quadrants for 200 ms. SOAs varied from 17 to 200 msec. When the mask followed the target with a short SOA, the face was often rendered invisible. After each trial, subjects indicated the location and the emotion of the target face with two button presses. In absence of an explicit visibility judgment, we coded trials where subjects responded correctly for *both* location and emotion as “visible”, and trials where subjects incorrectly identified the location as “invisible”. Summary measures of BM performance are shown in Figure S2b.

In the unmasked paradigms (UNM), stimuli were presented in a continuous train while subjects fixated the center of the display. Stimuli included upright faces, upside-down faces, houses, line drawings of tools, and Mondrian patterns used in CFS (but not flickering). Subjects indicated either the change of the fixation cross color (‘fixation task’) or, in separate experimental blocks, the repeat of stimulus category from trial to trial (‘one back task’). Since most trials did not require a response, UNM blocks were included for analysis if the subject made responses during the task, but no accuracy threshold was imposed. For the analyses in this paper, we did not distinguish between the fixation and one-back tasks.

## Computing Φ*

Φ* is a measure of integrated information in a candidate subsystem X, derived via a sequence of four computations (*I-IV*). For the detailed mathematical derivation of **Φ**\*, see (Oizumi, Amari, Yanagawa, Fujii, & Tsuchiya, 2015).

I. The states of a subsystem at time t, which we denote as *X*(*t*), is an *n*-dimensional vector, where its *i*^th^ dimension specifies the bipolar re-referenced voltage for the *i*^th^ channel. To estimate the uncertainty about the states of the mechanism, we employ the concept of entropy, under an assumption of Gaussian distribution of these states (Barrett & Seth, 2011; Oizumi, Amari, et al., 2015):

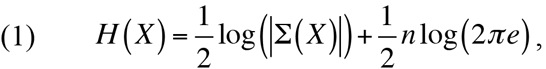

where Σ(*X* is the covariance matrix of X, estimated over the time interval [t-T/2, t+T/2] (T=200ms throughout the paper). The [*i*,*j*]^th^ element of Σ(*X* is the covariance between channel *i* and *j* of X over the time interval. |Σ(*X*)| is the determinant of Σ(*X*), known as the *generalized variance* (Barrett, Barnett, & Seth, 2010), as it describes the n-dimensional spread of instantaneous states of *X_t_*. The more different states *X* takes over the time interval, the more uncertain we are about its states at any time *t* within the interval.
II. Next, we estimate reduction in uncertainty about the mechanism’s states at *t* (=*X_t_*) given its past states (=X_t-t_, τ>0) using the concept of mutual information *I*:

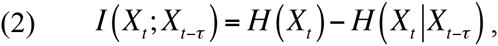

where *H*(*X_t_*|*X_t-τ_*) is the conditional entropy of the mechanism *X* across the delay *τ*. Under the Gaussian assumption, the conditional entropy is given by

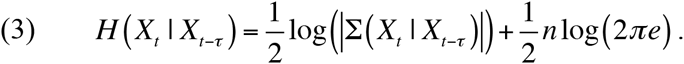 The covariance matrix of the conditional distribution, *X_t_*|*X_t-τ_* is given as

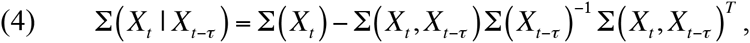

where Σ(*X_t_*,*X_t-τ_*) is the cross covariance matrix between *X_t_* and *X_t-τ_*, whose element Σ(*X_t_*, *X_t-τ_*) is given by covariance between i-th element of *X_t_* and j-th element of *X_t-τ_*. The way we use mutual information here is the same as auto-mutual information (Brenner, Strong, Koberle, Bialek, & Steveninck, 2000; Gómez, Hornero, Abásolo, Fernández, & Escudero, 2007; Julitta et al., 2011). *I(X_t_; X_t-_)* is a measure of the information that current states has about its *own* past states.
III. Integrated information Φ* over the subsystem *X* is information that cannot be partitioned into independent parts of *X* (For simplicity, we remove t and t- *τ* from X for the explanation of Φ* here). To identify integrated information in a subsystem, we estimate the portion of the mutual information that *can* be isolated in parts of X. The process consists of first defining the parts of *X* (a ‘partition’) and then estimating the fictitious mutual information assuming that the parts of a subsystem are independent. An estimate of ‘disconnected *I* is called mismatched information and denoted as *I*\* (Oizumi, Amari, et al., 2015; Oizumi et al., 2010). We compute *I*\* for *every possible partition* of *X*. For a subsystem of n channels, there are Bell(*n*)-1 partitions, where Bell(n) is the n-th Bell number (Bell, 1934). As an example, if *X* is made up of four ECoG channels, Bell(4) is 15, and thus there are 14 possible partitions of the subsystem (e.g. {a|bcd}, {ab|cd}, {ab|c|d},{a|b|c|d}, etc.), excluding the ‘trivial partition’ where all *n* elements are together in a single part (e.g. {abcd}). For each partition, we obtain Φ* = *I* - *I*\*. We select the partition *P* that minimizes the *normalized Φ*\* *value,* as defined in (Balduzzi & Tononi, 2008):

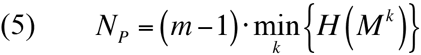

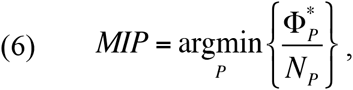

where m is the number of partitions and *M^k^* is the *k^th^* part of the system *X*. The normalization term *N_P_* counterbalances inevitable asymmetries introduced by computing Φ* across variable numbers of partitions of unequal size (Balduzzi & Tononi, 2008). The partition that minimizes normalized Φ* is called the ‘Minimum Information Partition’, or ‘MIP’. The MIP reveals the weakest link between the parts of *X*, the connections across which persists *only* information that is not reducible into parts.
IV. The integrated information of the subsystem is defined across the MIP as 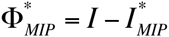(throughout this paper, Φ* refers to 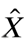. In our study, for stable estimation of covariance and cross-covariance, we used a shrinkage approach (Schäfer & Strimmer, 2005). By computing covariance and cross-covariance matrices separately for each trial and averaging these before computing the entropy, we estimated Φ* for bins of trial data. For the classification analyses, bins consisted of 3 trials each (randomly selected from a given percept/stimulus category); for the Φ* structure illustrations in Figures 3 and 4, all trials belonging to a particular stimulus/percept category were used to produce averaged covariance and cross-covariance matrices underlying the Φ* computations.

## Channel set selection procedure

We reasoned that where a stimulus evokes information integration, the structure of the integration should identify the percept, especially in visual areas. To test this hypothesis, we carried out the following procedure for each specific experimental setting (a particular stimulus task, as viewed through a particular brain region of electrode implantation - either ventral or lateral regions - in a particular subject). We identified the τ value and local set 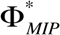 of four channels that contained the subsystem where the largest average Φ* was evoked (averaged over all 3-trial bins for all stimulus/percept categories):

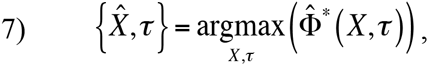

where 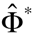 was the largest average *evoked* Φ* in a channel set *X* for a particular τ:

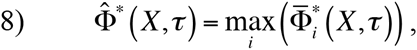

and the mean evoked 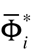 was the difference between the mean post-stimulus and mean pre-stimulus Φ* structures for the i-th subsystem within a channel set *X* for a particular τ:

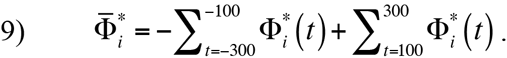

In 9), *t* is the center of a 200msec interval of ECoG data, and the Φ* estimates are averaged over all 3-trial bins for all stimulus/percept categories. Candidate channel sets included all positions of a rectangular searchlight in each subject’s lateral and ventral electrode arrays (a square searchlight for lateral arrays, and a slanted rhombus window for the ventral arrays). For the lateral arrays, most searchlight locations shared some 2^nd^-order subsystems with other overlapping locations, allowing some flexibility in the subsystem membership of the sampled Φ* structures; for the narrow ventral strips, the square searchlight shape did not allow any overlap, so the rhomboid shapes were added to allow for flexibility in subsystem membership. If the highest evoked Φ* value was located in more than one overlapping structure, the tie was resolved by the next highest evoked Φ* in the tied structures.

The ‘max evoked Φ*’ location 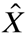 for each condition was included in the ANOVA and AUC timecourse (Figure 4b) described in the results. Evoked Φ* at each searchlight location (for clarity, excluding the closely overlapping ventral rhombuses), for each condition, is plotted in Figure S3. The ROI in Figures 1-3 was the max evoked Φ* system for S153’s ventral BM condition (the highest overall evoked Φ* over three stimulus paradigms). The max evoked Φ* system for the CFS condition was next to the depicted system as is clear from the figure.

Although Figure 4a shows the advantage of Φ* and MIP over I or H in most systems in the ventral and lateral cortex, using the above procedure to select ROIs for statistical analysis may seem to give AUC(Φ*) an unfair advantage over AUC(*I*) and AUC(*H*). In fact, AUC(Φ*) is correlated with evoked Φ*. To evaluate this problem, we also measured evoked *I* and *H* (both as ‘evoked maximum’ and ‘evoked minimum’, since entropy and mutual information tended to *decrease* after stimulus onset), and correlated all evoked measures with AUC, over all searchlight locations in all conditions. As shown in Figure S4A, while evoked I (and H) do correlate with AUC computed for I and H structures, they correlate as strongly or more strongly with AUC computed on Φ* structures. So, among the tested selection criteria (all of which, to some extent, are identifying regions of stimulus-evoked neural response), the highest AUC will tend to be obtained with Φ* structures. The Φ* AUC advantage even persists when we select single systems from each condition based on the minimal evoked *I* or *H* (Figure S4B,C) (rather than based on the maximal evoked Φ*, as in Figure 4b), although the overall level of classification tends to be poorer than when the criterion is based on Φ*.

## Multi-dimensional analyses for assessing the similarity of informational structures

In Figure 4, we used an unsupervised classification analyses for all available subjects within each stimulus paradigm and within recording locations (either lateral or ventral temporal surface), comparing the classification performance across the 4 measures (|Φ*|, MIP, I and H). As input features to the clustering, we reduced the dimensionality of the input features into the first 4 multidimensional scaling coordinates, and used the nearest-neighbor algorithm for percept classification (see below for details). |Φ*|, I and H coordinates were derived from (Pearson) correlation-distance matrices (Kriegeskorte et al., 2008); for a distance measure between MIP structures, we simply counted the proportion of subsystems with identical MIPs in the two compared structures.

Measurement of percept classification began with defining percept categories for each experimental stimulus interval. The percept categories were *Visible Face* and *Visible Mondrian* (including face-absent and masked-face intervals) for CFS, *Visible Face* and *Visible Noise* (including masked-face and face-absent ‘catch’ trials) for BM; for UNM there were four possible percepts: *Faces* (including upright and inverted faces), *Houses, Mondrians,* and *Tools*). For BM and CFS data, the assignment of trial intervals to percept categories was based on psychophysical responses as described in the earlier section; so, ‘face’ bins were labelled according to visibility judgments and objective accuracy.

We assessed the clustering with a cross-validation procedure. First, using 70% of randomly sampled trials within each percept category as a training set, we determined the center of gravity for each category. Second, using the remaining 30% of trials as a test set, we computed receiver-operating characteristic (ROC) curves for each class by gradually extending a criterion distance from the category center; we then averaged the area under the curve (AUC) over all categories as the measure of the classification accuracy. We varied the criterion radius for each category from the mean estimated from the training set, counting the number of same-category bins within the radius as ‘Hit’ and other-category bins as ‘False Alarm’. By smoothly varying the radius from 0 to infinity, the proportion of both Hit and False Alarm changes from 0 to 1. ROC curves connect the dots from [0,0] to [1,1]. The procedure was repeated 20 times, with random resampling of train/test bins. For statistical analysis, each AUC value was converted into z-scores.

## Author contributions

AMH and NT conceived the concepts of the study and conducted the analyses. CK, HK, MAH, RA and NT conceived the experimental study. HK, HO, and MAH performed neurosurgery. CK, HK, and NT collected the data. HK, HO and NT carried out preliminary analyses including anatomical localization of electrodes. MO invented the Φ* measure, provided relevant code, and discussed critical aspects of the analyses. AMH and NT wrote the paper. All the authors read and commented on the manuscript.

## Acknowledgements

AMH was supported by an Endeavor Fellowship from the Australian Department of Education. CK, HK, HO, and MAH were supported by NIH grants R01 DC004290 and UL1RR024979. NT was supported by the Future Fellowship (FT120100619) and the Discovery Project (DP130100194) from the Australian Research Council. We thank all subjects for their participation in the study, the medical team at University of Iowa Hospital for their assistance, Haiming Chen for his technical assistance and Jochem van Kempen and Fabiano Baroni for their parallel analyses of percept/stimulus decoding in the same dataset.

---

**Figure S1.**
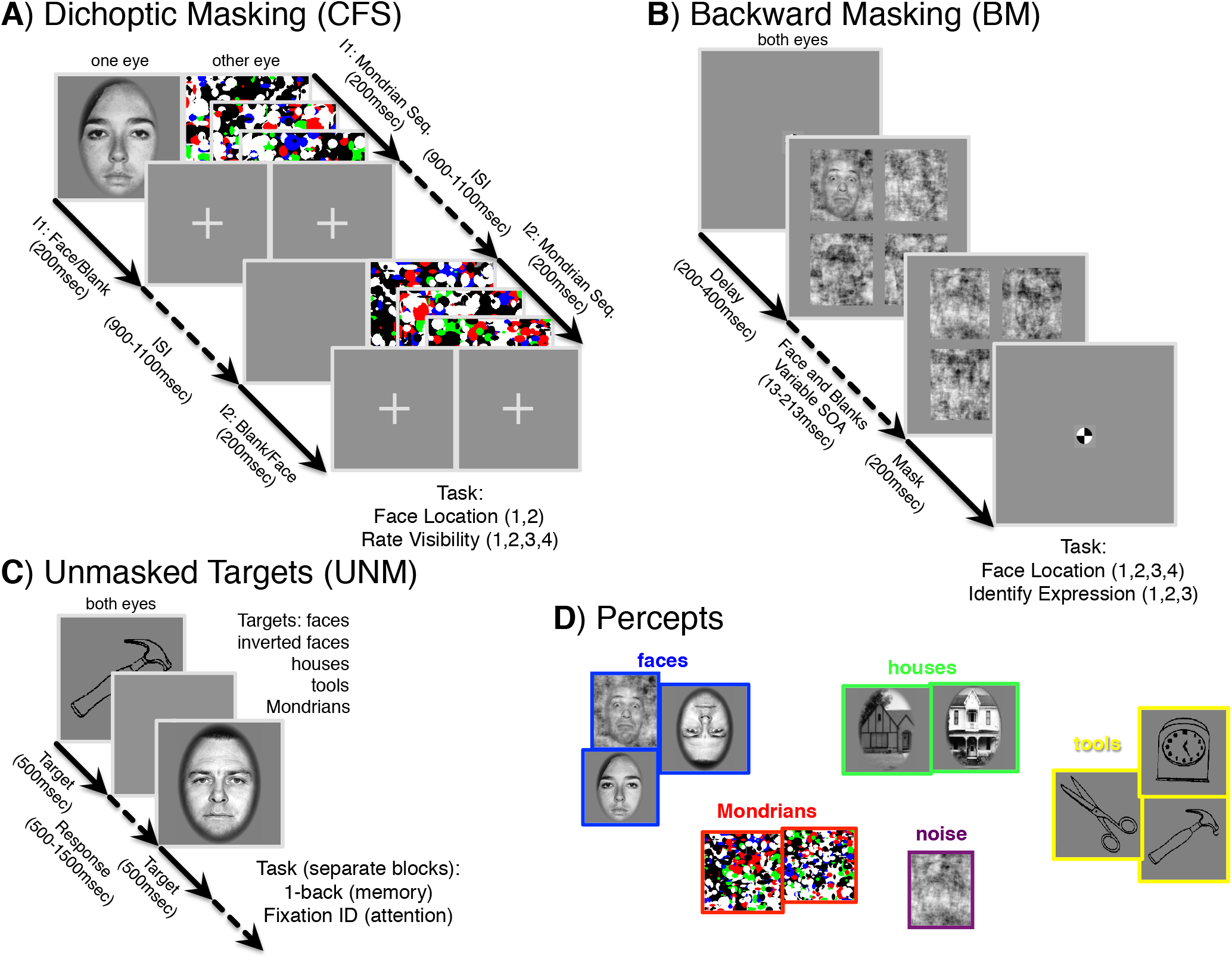
Paradigms for stimulus presentation. **A**. Continuous flash suppression (CFS) task. Each trial consisted of a temporal sequence of two stimulus intervals, separated by a random inter stimulus interval (ISI, 900-1100ms). In one eye, each interval contained three flashes of colorful Mondrian patterns; in the other eye, *one* interval contained a face image of variable contrast. These conditions result in stochastic trial-to-trial visibility of the target face; sometimes the face is consciously seen, sometimes it is not. The subjects’ task was to select which interval contained the face, and to indicate how visible it was on a scale of 1 to 4. **B**. Backward masking (BM) task. After a random fixation delay, subjects saw an array of four noise patches, one of which contained a face image (the upper left, in the illustration), for 13 msec. After a variable stimulus onset asynchrony (SOA), another array of noise patches was presented to reduce the visibility of the face. The subject’s task was to identify the location of the face target among 4 possible locations, and also to identify its emotional expression among 3 possible labels (happy, fearful and neutral). C. Unmasked conditions included a one-back memory task, in which subjects paid attention to the category of the stimuli, or a fixation task, in which they ignored the category of the stimuli. In both tasks, faces and other objects were presented for 500 msec without any masks, with trials separated by a blank interval (500ms for the fixation task, 1000 msec for the one-back task). **D**. From subjects’ performance on a task (correct/incorrect, ratings of visibility, identification of expression), we can reasonably infer what their percept was likely on each trial of an experiment. We used subjects’ performance to divide trials into the percept categories shown here: faces (and inverted faces), houses, tools, Mondrians, and noise.

**Figure S2.**
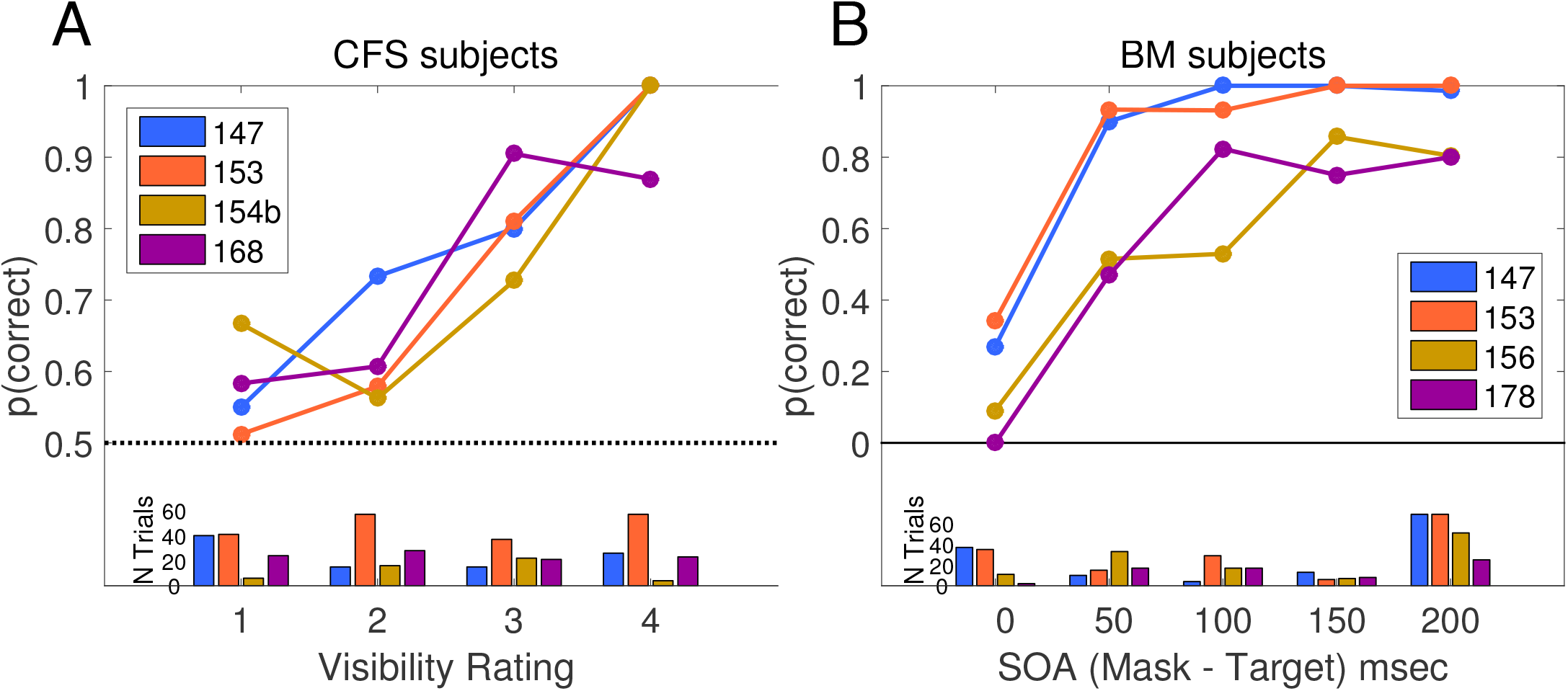
Behavioral performance on the masking tasks, by two groups of four patients (indicated by colors/number IDs). Upper panels show proportion correct for respective targets in CFS (A) and BM (B) tasks. Lower panels show the number of trials at each trial type. A) Proportion correct for four CFS subjects, as a function of visibility rating. Trials rated as ‘3’ or ‘4’ were treated as ‘visible’ in the classification analyses. B) Proportion correct for BM subjects, as a function of (binned) backward-mask SOA. Trials where subjects were correct on both 4AFC location and 3AFC emotion judgments (here coded as ‘correct’) were treated as ‘visible’ in the classification analyses.

**Figure S3 (in two parts).**
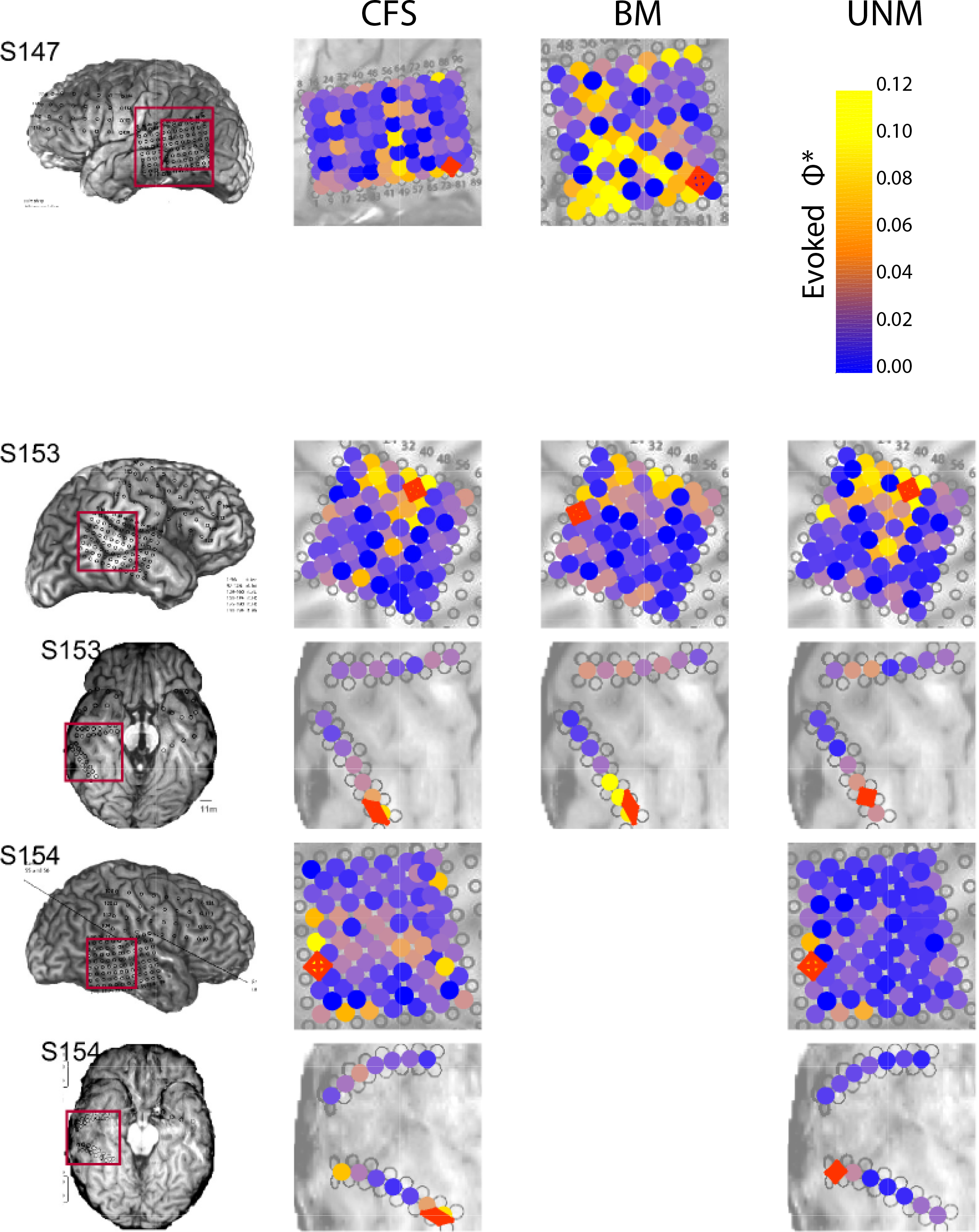
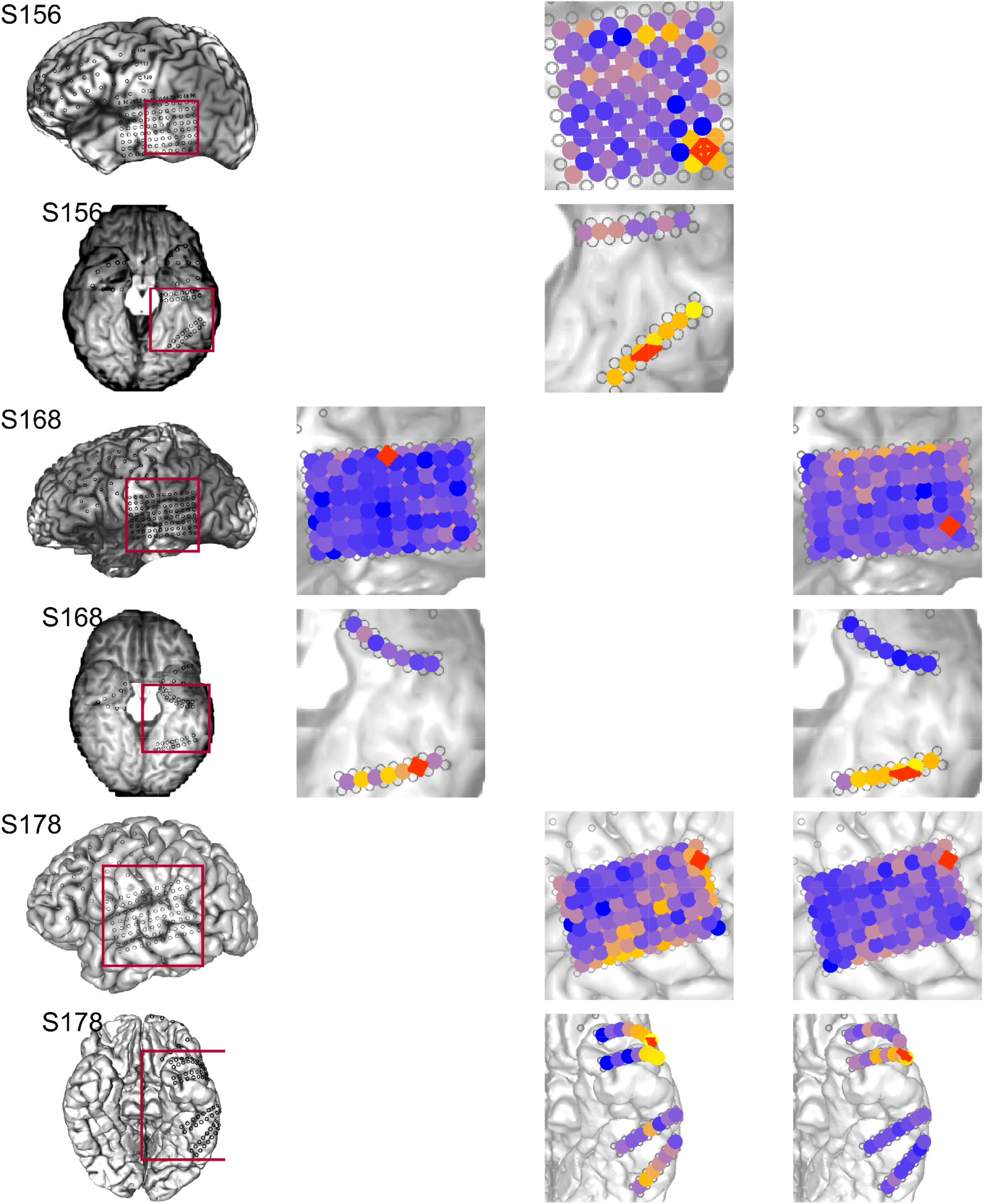
Max evoked Φ* searchlight and ROI selection. For each of 22 conditions (combinations of 6 subjects with ventral and/or lateral electrode implantation, and 3 stimulus paradigms, as represented by the rows and columns of the figure), we computed the average Φ* structure (over all experiment trials) for every ‘closed’ 4-channel system (square and elongated rhombuses where each vertex is adjacent to the next), at each of four x values (1.5, 3, 6, and 12 msec). As an ROI for further analysis, we chose one channel set that contained the subsystem with the highest Φ* regardless of stimulus/percept (see Methods for details). The selected regions for classification analysis (Figures 1, 2, 3, and 4b) are marked in red. Marker colors encode the maximal evoked Φ* at the centroid of each system (values above 0.1 are all given the color yellow).

**Figure S4.**
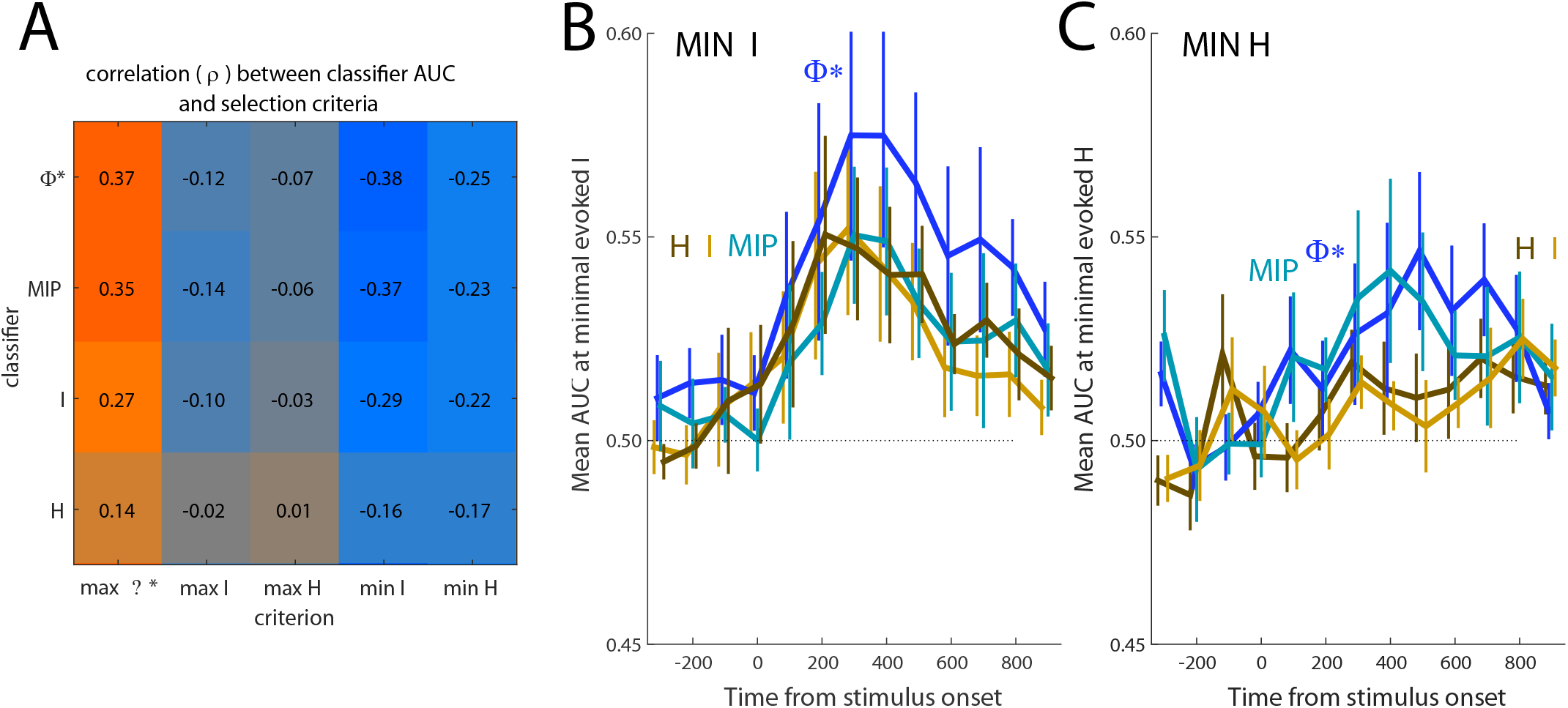
**A**) Correlation over all searchlight locations of classification accuracy AUC for different information structures (Φ*, MIP, I, and H) with different structure properties (maximum evoked Φ*, maximum evoked I or H, or minimum evoked I or H). For the main analyses in the paper, we selected cortical regions where we observed the largest average increase in Φ* in response to *all* stimulus conditions. Evoked Φ* correlates with AUC, most strongly with AUC(Φ*) and AUC(MIP), reflecting the result shown in Figure 4b. However, Φ* advantage is not due to the selection criterion; minimum evoked I or H both correlate with classification accuracy - but the correlation is still best for Φ* and MIP structures, meaning that if we choose a region for analysis based on the greatest *decrease* in mutual information, the best classifier in that region will still tend to be the Φ* structure, not the I (or H) structure. B,C) Selecting systems based on a “minimal mutual information or entropy” criterion still picks out systems where Φ* is a better classifier, although peak AUC is not as high as when the criterion is max evoked Φ* (as in Figure 4b).

**Video 1**: 4-channel Φ* structure (from Figures 2 and 3), rotated through three dimensions to clearly illustrate its construction. The x-y coordinates are arranged for aesthetic purposes, to illustrate the layout of the Hasse graph that connects all the subsystems. The z-coordinate is the magnitude of Φ* for each subsystem. This structure is generated by a system of four channels over subject 153’s posterior fusiform gyrus.

**Video 2**: 4-channel Φ* structures for visible BM faces (left structure), visible CFS faces (middle structure), and invisible CFS faces or Mondrians (right structure). Similar structures from the same subject 153, are shown in Figure 4.

